# Using methyl bromide for interspecies cell-cell signaling and as a reporter in a model soil consortium

**DOI:** 10.1101/2023.09.06.556470

**Authors:** Jiwoo Kim, Li Chieh Lu, Xiaodong Gao, Kirsten S. Hofmockel, Caroline A. Masiello, Jonathan J. Silberg

## Abstract

Soil microbial communities with reduced complexity are emerging as model systems for studying consortia-scale phenotypes. To establish synthetic biology tools for studying these communities in hard-to-image environmental materials, we evaluated whether a single member of a model soil consortium (MSC) can be programmed to report on gene expression without requiring matrix disruption. For these studies, we targeted a five member MSC that includes *Dyadobacter fermentans*, *Ensifer adhaerens*, *Rhodococcus* sp003130705*, Streptomyces* sp001905665, and *Variovorax beijingensis*. By coupling the expression of a methyl halide transferase to a constitutive promoter, we show that *Variovorax beijingensis* can be programmed to synthesize methyl halides that accumulate in the soil headspace at levels that are ≥24-fold higher than all other MSC members across a range of environmentally-relevant hydration conditions. We find that methyl halide production can report on a MSC promoter that is activated by changes in water potential, and we demonstrate that a synthetic gas signal can be read out directly using gas chromatography and indirectly using a soil-derived *Methylorubrum* that is programmed to produce a visual output in response to methyl halides. These tools will be useful for future studies that investigate how MSC respond to dynamic hydration conditions, such as drought and flood events induced by climate change, which can alter soil water potential and induce the release of stored carbon.

## Introduction

Soils are complex living materials whose health and productivity are critical to the biosphere because of their roles in providing an environment for plants, filtering and cleaning water, and regulating carbon storage.^1–3^ Soil harbors tremendous biological diversity,^4^ communities of microbes whose collective metabolic activities and phenotypes are responsible for soil functions.^5^ While -omics approaches can be applied to bulk soil samples to provide high-resolution information about who is present and what are they doing at any point in time,^6, 7^ such as consortium-scale gene and protein expression, we cannot yet predict how the metabolic activities of individual soil consortium members dynamically change as environmental resources vary.^8^ In part, this challenge exists because -omics tools are disruptive to soils and only provide snapshots of microbial behaviors, rather than dynamic information.^9^ Currently, there is a need to understand how dynamic changes in environmental resources affect consortia-scale phenotypes, termed the metaphenome.^10^ To make this problem tractable, model soil communities have recently been developed to enable studies of population dynamics and cell-cell interactions mediated by metabolites and signals.^7, 11^

Synthetic biology is increasingly being explored as a strategy to alter soil biology and improve soil health and productivity. Because of the importance of root microbiota for agricultural yields, trans-kingdom plant-to-microbe signaling has been developed as a strategy to control gene expression in rhizosphere bacteria.^12^ Rhizosphere bacteria have been programmed to visually respond to a synthetic signal, and this approach has been used to engineer control over nitrogen fixation.^13^ Synthetic biology is also being applied in bulk soils to enhance microbial bioremediation of contaminants, *e.g.*, pesticides,^14, 15^ and to understand soil processes outside of the rhizosphere.^16, 17^ To achieve the latter, soil biosensors have been developed that report on gene expression non-disruptively by producing indicator gases, such as methyl halides and ethylene.^18, 19^ To date, these volatile metabolites have been used to report on the presence of synthetic microbes in soil,^20^ the effects of hydration on horizontal gene transfer,^18^ and the effect of the soil matrix on signal bioavailability.^19, 21^ While gas signals can be measured in the headspace of soils containing synthetic cells at titers as low as ∼10^3^ microbes per gram of soil,^20^ it remains unclear if they can be applied in soils to control the flow of information about individual microbial behaviors.

A model soil consortium (MSC) was recently developed by enriching a community of interacting soil microbes and showing that they can be stored, revived, and shared.^7^ While this MSC represents an opportunity to study how changes in the physicochemical properties of a soil affect microbe-microbe interactions and the metaphenome, we lack simple synthetic biology tools to monitor the behaviors of individual microbes in this MSC. To overcome this challenge, we show that one member of this MSC can be programmed to generate a methyl halide signal that greatly exceeds the background methyl halide generated by other MSC members, and we demonstrate the utility of this approach by characterizing an MSC promoter in both a liquid and soil environment. We find that this synthetic signal can be read out directly using gas chromatography mass spectrometry (GC-MS) or indirectly using a *Methylorubrum extorquens* CM4 biosensor,^22^ a soil-derived microbe that expresses a fluorescent protein in response to methyl halide sensing. Furthermore, using soil habitats that range in size from 1 to 50 grams of soil, we show that methyl halide cell-cell signaling can be achieved in soils hydrated across a range of levels observed in the environment. These tools are expected to enable new types of metaphenome studies in hard-to-image living environmental materials.

## Results and Discussion

### Gas reporting in a soil consortium

MSCs of varying complexity have been identified that can be stored and revived for soil studies.^7, 11^ These consortia represent an opportunity to advance soil biosensors by testing the compatibility of native soil microbes with indicator gas reporters.^18–20^ To investigate whether these reporters are compatible with MSC studies, we first evaluated whether individual MSC members present a gas signal that can be detected using standard GC-MS analysis. These measurements focused on five microbes from a chitin-degrading consortium of microbes,^7^ including *Dyadobacter fermentans* (*Df*), *Ensifer adhaerens* (*Ea*), *Rhodococcus* sp003130705 (*Rh*), *Streptomyces* sp001905665 (*St*), and *Variovorax beijingensis* (*Vb*).

For each microbe, gas production was evaluated in R2A growth medium containing 10 or 100 mM NaBr (Figure 1A), which represents halide levels that exceed the concentrations observed in many soil settings.^23^ Halides were varied as this is an essential substrate for Methyl Halide Transferase (MHT) indicator gas reporters.^24^ As a frame of reference, we evaluated indicator gas produced by an *E. coli* strain (*Ec-mht*) that has been engineered to constitutively express *Batis maritima* MHT.^18^ With these measurements, methyl bromide was detected in the headspace of every culture following a 48 hour incubation (Figure S1). When this gas production was normalized to the optical density (OD) of each culture to account for differences in cell number (Figure 1B), we found that the indicator gas signal from each MSC member was more than two orders of magnitude lower than *Ec*-mht. With both NaBr concentrations, *Df* presented the highest CH_3_Br signal, which was 562-fold lower than the *Ec*-*mht* signal in the presence of 10 mM NaBr and 370-fold lower than the *Ec*-mht signal in the presence of 100 mM NaBr. A similar trend was observed when the CH_3_Br signal was normalized to respiration (Figure 1C), a strategy used to quantify indicator gas signals in soils.^25^ These results show that MSC members synthesize CH_3_Br when grown in the presence of high, non-physiological levels of NaBr, albeit at levels that are more than two orders of magnitude lower than the concentration that is produced by a synthetic microbe.

**Figure 1.**
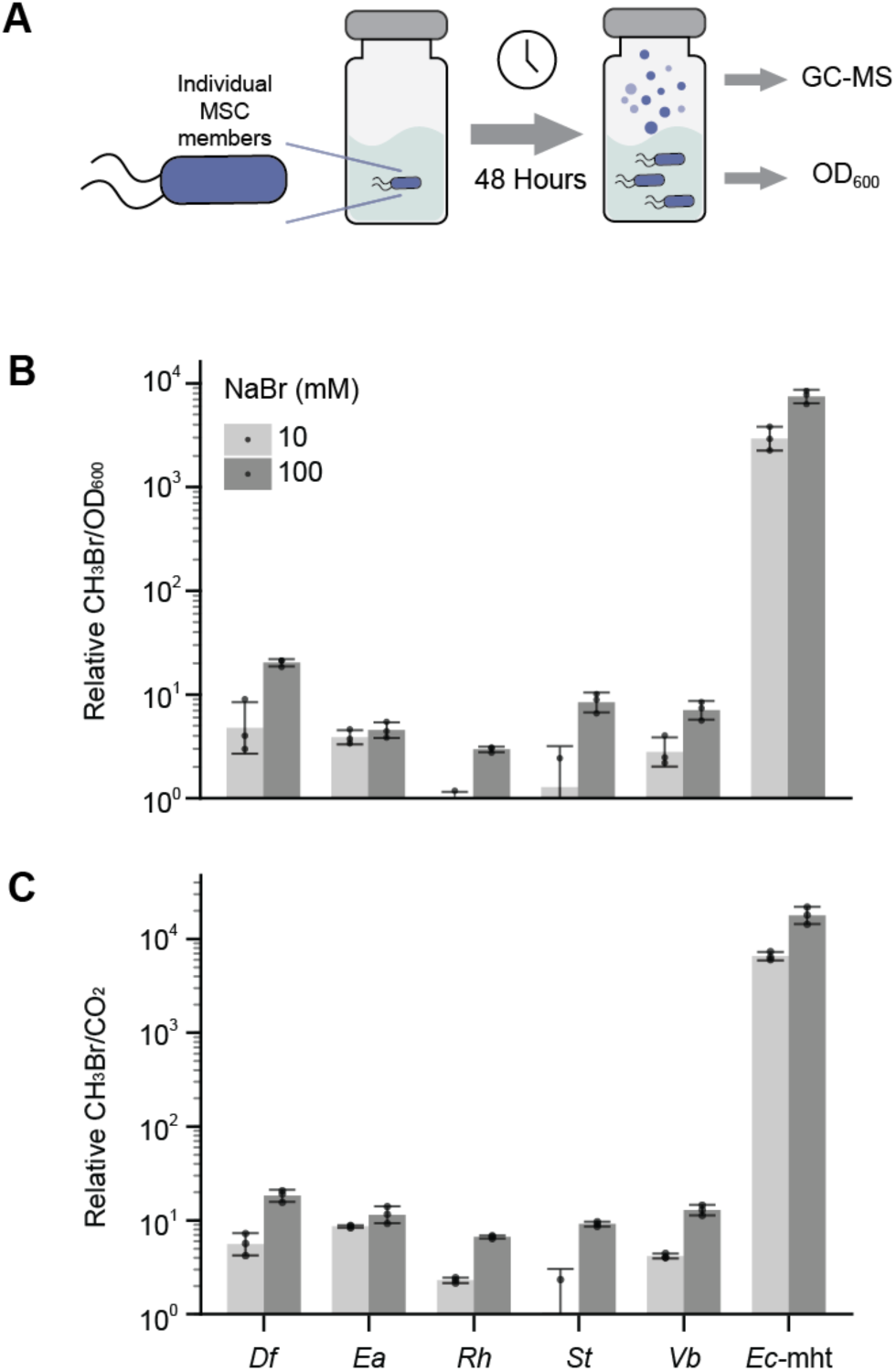
MSC members produce low but detectable CH_3_Br. (**A**) Scheme for measuring MSC CH_3_Br production in liquid culture. Cells (1 mL, OD = 0.05) in vials (2 mL) containing R2A medium were capped and incubated for 48 hours at 30°C, and the headspace gas was measured. Gas production for individual MSC isolates normalized to (**B**) OD and (**C**) respiration. These experiments were performed in medium supplemented with 10 mM (light gray) or 100 mM (dark gray) NaBr. All MSC members presented a signal that was significantly lower (p < 0.01; two-tailed, unpaired, t-test) than an *E. coli* strain that constitutively expresses an MHT (*Ec*-mht), including: *D. fermentans* (*Df*), *E. adhaerens* (*Ea*), *R.* sp003130705 (*Rh*), *S.* sp001905665 (*St*), and *V. beijingensis* (*Vb*). Error bars represent ±1 standard deviation from three biological replicates, and all data is normalized to the lowest signal which was given a value of one.

We hypothesized that a single MSC member could be programmed to synthesize a unique CH_3_Br signal that exceeds the background levels produced by the other MSC members by introducing a plasmid that constitutively expresses an MHT, since these enzymes require a common metabolite, S-adenosyl methionine, and a halide ion as substrates.^24^ We chose to target *Vb* because it is easy to grow and transform with plasmids using conjugation, and microbes from the genus have been shown to have beneficial interactions with other bacteria and plants and to have metabolic features to degrade common pollutants found in soil and water.^26^ As a frame of reference, we evaluated *E. coli* gas production using the same vector, since this strain has been applied extensively as a gas-reporting biosensor.^18^ *Vb* transformed with this vector (*Vb-mht*) presented a robust CH_3_Br signal across three different growth conditions (Figure S2), including rich medium (R2A), minimal medium (M63), and a diluted minimal medium (MIDV1) whose osmolarity more closely reflects that in soils.^25^ When the CH_3_Br signal was normalized to respiration (Figure 2A), the *Vb-mht* signal in R2A medium was similar to that generated by *E. coli* harboring the same vector, *Ec-mht*, while it was significantly higher in R2A and M63 medium. With wildtype *E. coli*, the signal was below the limit of detection as previously reported,^18^ while *Vb* presented a detectable signal in a subset of the conditions. These results show that a single microbe from the soil-derived MSC can be programmed to present enhanced indicator gas production across a range of growth conditions.

**Figure 2.**
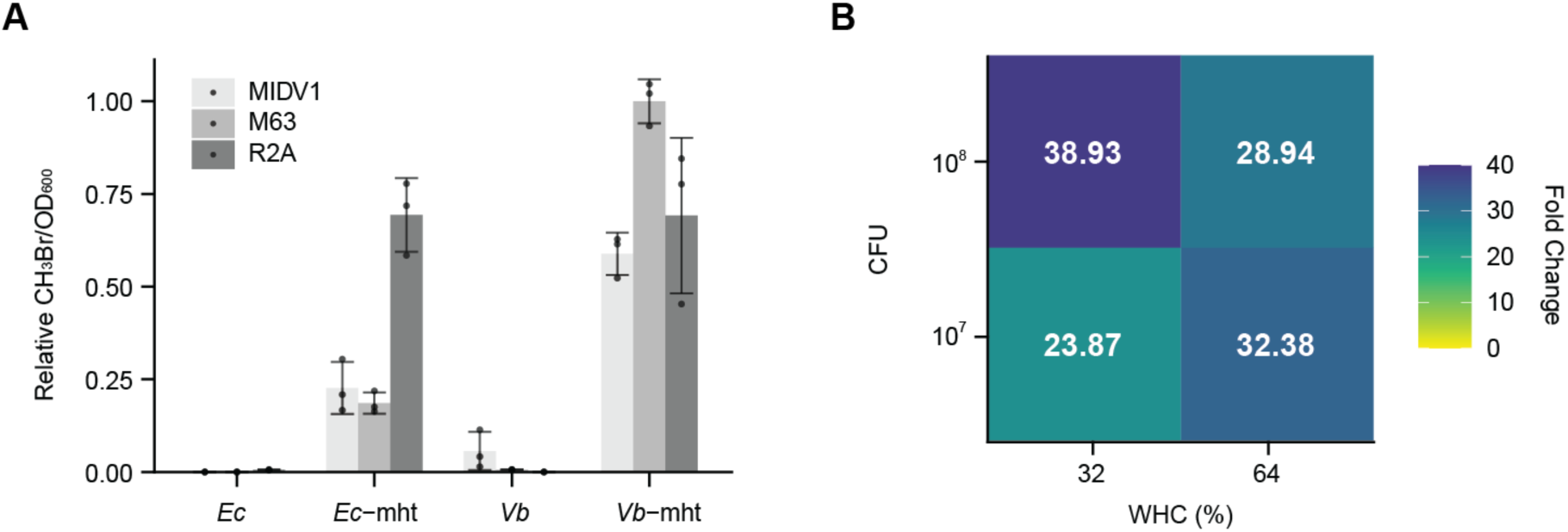
Programming gas reporting in an MSC microbe. (**A**) When transformed with a vector that constitutively expresses an MHT, *E. coli* (*Ec*-mht) and *Vb* (*Vb*-mht) present significantly higher gas production than native cells in different growth media, including R2A (black), M63 (white), and MIDV1 (gray) medium containing 100 mM NaBr (\**p* < 0.01; two-tailed, unpaired, t-test). (**B**) The ratio of the signal obtained from *Vb*-mht/*Vb* in soil containing different cell titers (10^7^ or 10^8^ CFUs), hydrated to different environmentally-relevant water holding capacities (WHC). Across all conditions, MHT-expressing cells produce a significantly higher gas signal (CH_3_Br/CO_2_) than cells lacking an MHT (\**p* < 0.05; two-tailed, unpaired, t-test). Values shown represent the mean from three biological replicates ±1 standard deviation.

To investigate if soil affects the *Vb* gas signal, we evaluated the accumulation of CH_3_Br in the headspace of a soil containing different titers of *Vb-mht* (10^7^ or 10^8^ CFU) held at different hydration levels (32% or 64% field capacity). For these experiments we used a Texas Alfisol, which we previously characterized.^20^ Soils were hydrated using MIDV1 containing 100 mM NaBr, because this medium is designed to have an osmotic pressure closer to the range found in natural soil conditions.^25^ As observed in liquid medium, native *Vb* generated low levels of headspace CH_3_Br when added to the Texas Alfisol (Figure S3). When the headspace CH_3_Br signal was normalized to CO_2_, which was done to account for variation in titer and metabolism, the normalized signal increased significantly across all conditions analyzed, increasing 24 to 39-fold (Figure 2B). These results show that *Vb* can be programmed to report on gene expression non-disruptively in a soil held at environmentally-relevant hydration levels.

### Characterizing an MSC promoter in soil

To investigate whether our gas reporter can be used to characterize an MSC promoter in soil, we created a vector that expresses MHT using a promoter from *Vb* coding sequence CDS.21, which is predicted to be hyperosmotic responsive.^27^ We chose this promoter because soil microbes experience osmotic challenges as environmental conditions cycle between wet and dry.^28^ While osmotic responsive promoters are often characterized by adding high solute concentrations to increase the osmotic potential in liquid cultures,^29, 30^ it is possible that matric potential of a soil could also activate these promoters.^31, 32^ The latter induction is challenging to measure using bulk soils using biosensors, as soils are hard-to-image.^33^ Real-time, *in situ* detection of microbial responses to osmotic changes is becoming increasingly important for understanding microbial responses to climate change, including drought and flood events.^34, 35^

We first evaluated the effect of osmotic potential on the growth of *Vb* by growing cells in MIDV1 cultures containing 0 to 600 mM sucrose, a carbon source that *Vb* is unable to use to make biomass.^7^ This range of sucrose concentrations generates osmotic pressures that range from -319 to -1807 kPa; the latter is higher than the permanent wilting point for many plants.^36^ This analysis revealed that cell growth is decreased when sucrose concentrations are ≥400 mM and osmotic potential is lower than -1300 kPa (Figure S4). To investigate whether the CDS.21 promoter is regulated by osmotic potential, we evaluated the effect of a similar range of sucrose concentrations on gas production from *Vb* that uses this promoter to express the MHT, *Vb-21-mht*, and cells containing an empty vector. As sucrose increased from 0 to 400 mM, CH_3_Br production increased with both cell types, while higher levels of sucrose decreased CH_3_Br levels. When the gas production was normalized to OD, only *Vb-21-mht* presented sucrose-induced gas production (Figure 3A). When we calculated the ratio of the indicator gas signal from *Vb-21-mht* to cells containing an empty vector, we observed ratios that ranged from 2.4 (at -319 kPa) to 11.6 (at -1807 kPa). These results show that the CDS.21 promoter is induced by osmotic stress, and they calibrate the pressures that induce transcription.

**Figure 3.**
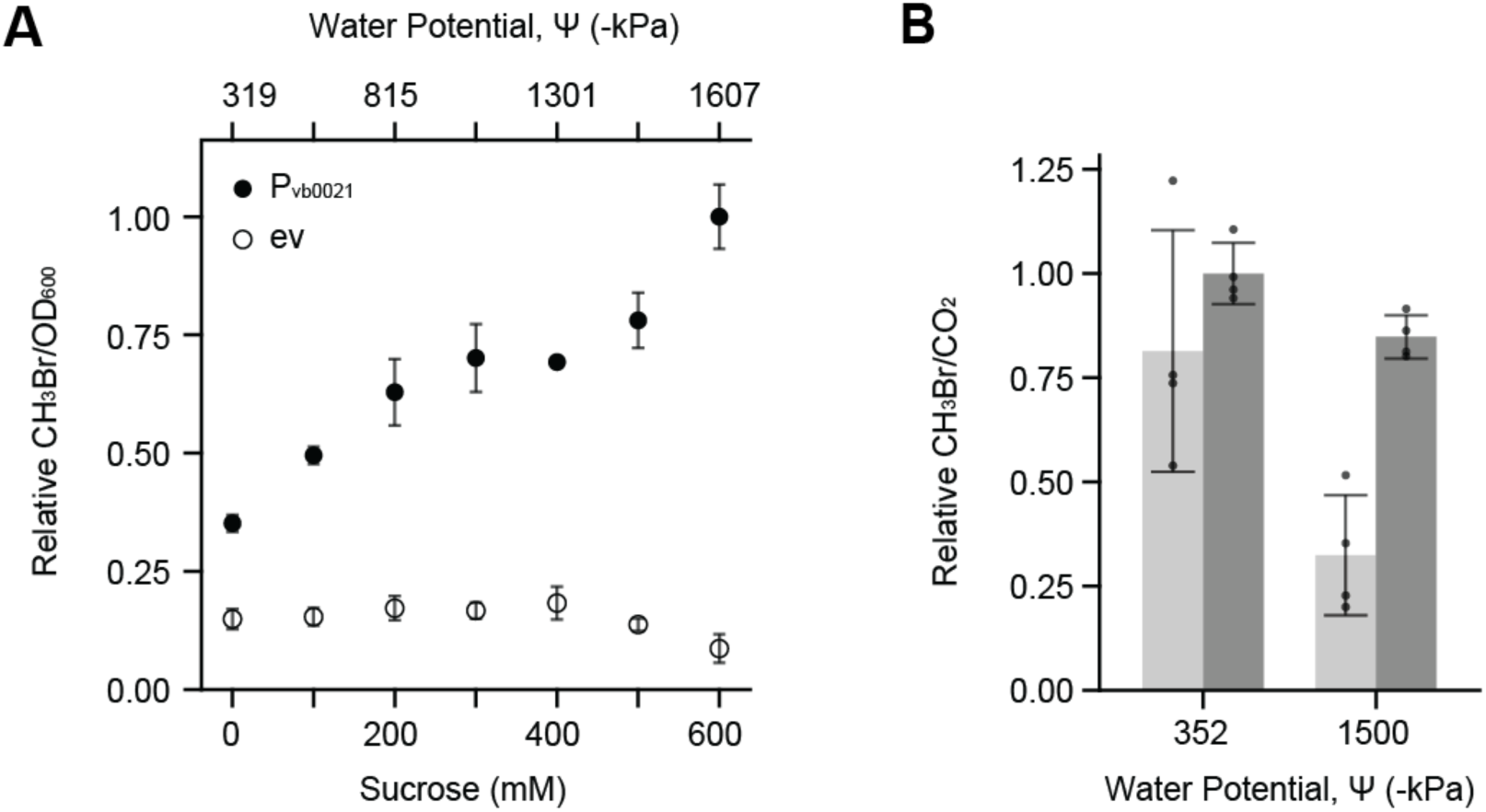
Using indicator gas reporting to monitor the effects of osmotic potentials on a native *Vb* promoter. (**A**) Headspace CH_3_Br generated by *Vb* cultures containing an empty vector or a vector that uses the *Vb* promoter *P_vb0021_* to regulate MHT expression (filled circles). *Vb* transformed with each plasmid was grown in MIDV1 containing NaBr (20 mM) and varying concentrations of sucrose, which alters the water potential of the culture. Cells harboring the vector that uses *P_vb0021_* to regulate MHT expression presented a significant increase in signal with sucrose addition compared with cells grown in the absence of sucrose (***p < 0.0005, two-tailed, unpaired t-test). Each point plotted represents the mean of 4 or more biological replicates and error bars represent ±1 standard deviation. (**B**) Each cell type was added to soil hydrated to 100% water holding capacity using MIDV1. When soil was hydrated using MIDV1 lacking sucrose (-352 kPa) the signal from the cells that use *P_vb0021_* to regulate MHT expression (dark gray) could not be differentiated from those containing empty vector (light gray). However, the signal from the former was significantly higher in soil containing sucrose (463 mM) hydrated to the same level, which results in a water potential of -1500 kPa (***, *p* = 0.0005, two-tailed, unpaired, t-test). The bars represent the arithmetic mean of 4 biological replicates, with error bars shown that represent ±1 standard deviation, while the value of each replicate is shown as a black circle within the bar. All data is scaled relative to this highest indicator gas signal, which was given a value of one.

In soil, the total water potential is expected to differ from liquid cultures, as cells experience both osmotic and matric potential.^37^ To investigate whether the CDS.21 promoter responds to changes in osmotic potential in a matrix environment where the total potential is determined by the osmolytes and matrix, we evaluated gas production in a synthetic soil (Q3a) whose matrix potential was previously described.^25^ When this artificial soil was hydrated to field capacity with MIDV1 (-352 kPa), *Vb-21-mht* presented an indicator gas signal that was similar to cells containing an empty vector (Figure 3B, Figure S5). When sucrose was added to the soil to increase the total potential to -1500 kPa, the *Vb-21-mht* signal was significantly higher than cells containing an empty vector (\*\*\**p* = 0.0005; two-tailed, unpaired, t-test). These results show that a MHT reporter can be used to monitor how osmotic and matric potential work together to regulate promoter activities.

### Calibrating a methyl halide biosensor

*M. extorquens* CM4 is a soil-derived methyl halide degrading bacterium that can use this one carbon molecule as a source of carbon and energy.^38^ Transcriptional studies have shown that this microbe has numerous promoters that are activated by the methyl halides,^39^ and one of these has been used to create a methyl halide biosensor.^22^ To investigate if this latter strain can be used to program interspecies methyl halide signaling, we performed an experiment in which headspace gas from an *Ec-mht* culture was injected into the headspace of a culture containing an *M. extorquens* CM4 *receiver strain* (Figure 4A), which was programmed to regulate the expression of yellow fluorescent protein (YFP) using the cmuA promoter *P_cmuA_*. For these experiments, *Ec-mht* cultures were grown to stationary phase in sealed vials, and different volumes of headspace gas were injected into sealed vials containing the *M. extorquens* CM4. As a control, headspace gas from native *E. coli* cultures was transferred to *M. extorquens* CM4 vials. With these experiments, the whole cell fluorescence of *M. extorquens* CM4 increased as increasing volume of headspace gas was injected from *Ec-mht* cultures but not *E. coli* cultures (Figure S6). When the fluorescence signal was normalized to the optical density of biosensor cultures (Figure 4B), a significant increase in whole cell fluorescence was observed in *M. extorquens* CM4 cultures containing ≥10 µL of gas from the *E. coli* producer cultures. In contrast, injection of headspace gas from native *E. coli* cultures had no effect on *M. extorquens* CM4 fluorescence. These findings show that gas-producing microbes generate sufficient headspace gas to activate gene expression in *M. extorquens* CM4.

**Figure 4.**
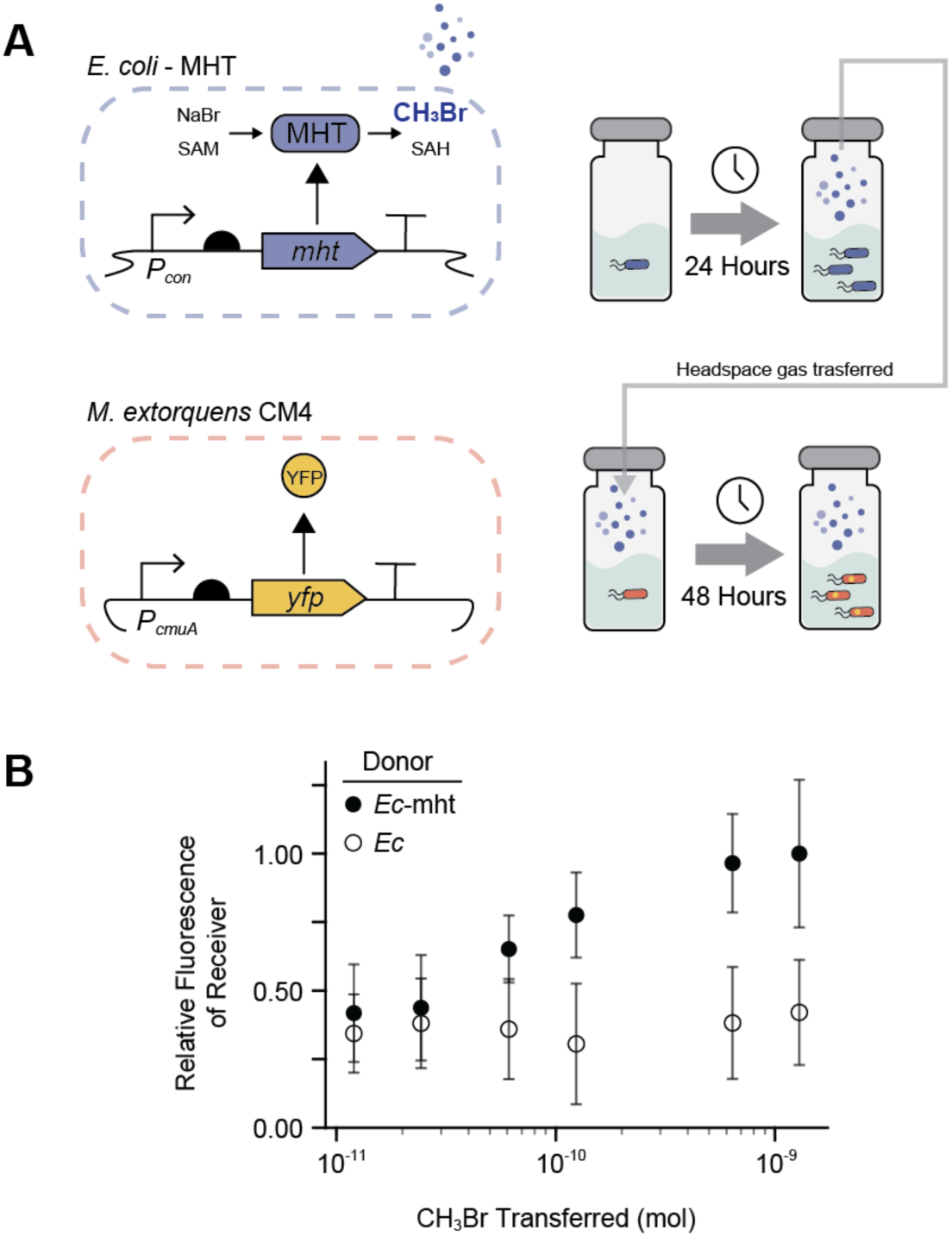
*M. extorquens* can report on biological indicator gas production. (**A**) Genetic circuits used to program (*top*) CH_3_Br production in an *E. coli* sender strain and (*bottom*) CH_3_Br detection using a *M. extorquens* CM4 receiver strain. Following a short incubation of the sender strain (blue). in a closed system, different volumes of headspace gas were transferred to closed systems containing the receiver strain (yellow), which were subsequently incubated prior to measuring their absorbance and fluorescence. (**B**) Receiver strains exposed to headspace gas from *Ec*-mht cultures (closed circles) presented significantly higher signal than strains exposed to headspace gas from native *E. coli* in cases when the headspace CH_3_Br after transfer was >167 nM (*p* < 0.05; two-tailed, unpaired, t-test). Relative fluorescence values are normalized to the maximum signal. Data represent the mean of seven biological replicates and error bars represent ±1 standard deviation.

We next investigated whether the headspace gas from the MSC culture grown in the presence of 10 or 100 mM NaBr contains sufficient methyl halides to activate gene expression in *M. extorquens* CM4. For these experiments, the five different MSC microbes were grown as a mixture in R2A liquid medium for two days, prior to injecting headspace gas into sealed vials containing the methyl halide biosensor. We also performed parallel experiments where *Vb-mht* was included in the MSC rather than native *Vb*. The CO_2_ levels produced by the MSC containing *Vb* or *Vb-mht* was similar across all experiments (Figures S7A-B), while the CH_3_Br signal was significantly higher in the headspace of MSC cultures containing *Vb-mht*. Also, the normalized headspace CH_3_Br/CO_2_ signal was ≥26-fold higher in the MSC cultures containing *Vb*-mht cultures compared to those MSC cultures containing native *Vb* (Figure 5A). The total headspace CH_3_Br varied with halide supplementation, with MSC cultures containing 100 mM NaBr presenting 3-fold more indicator gas in the headspace compared to cultures containing 10 mM NaBr, as did the ability of this headspace gas to activate YFP production within *M. extorquens* CM4 (Figure 5B, Figures S7C-D). These results show that one member of the MSC can be programmed to produce indicator gas at a level that activates gene expression in *M. extorquens* CM4.

**Figure 5.**
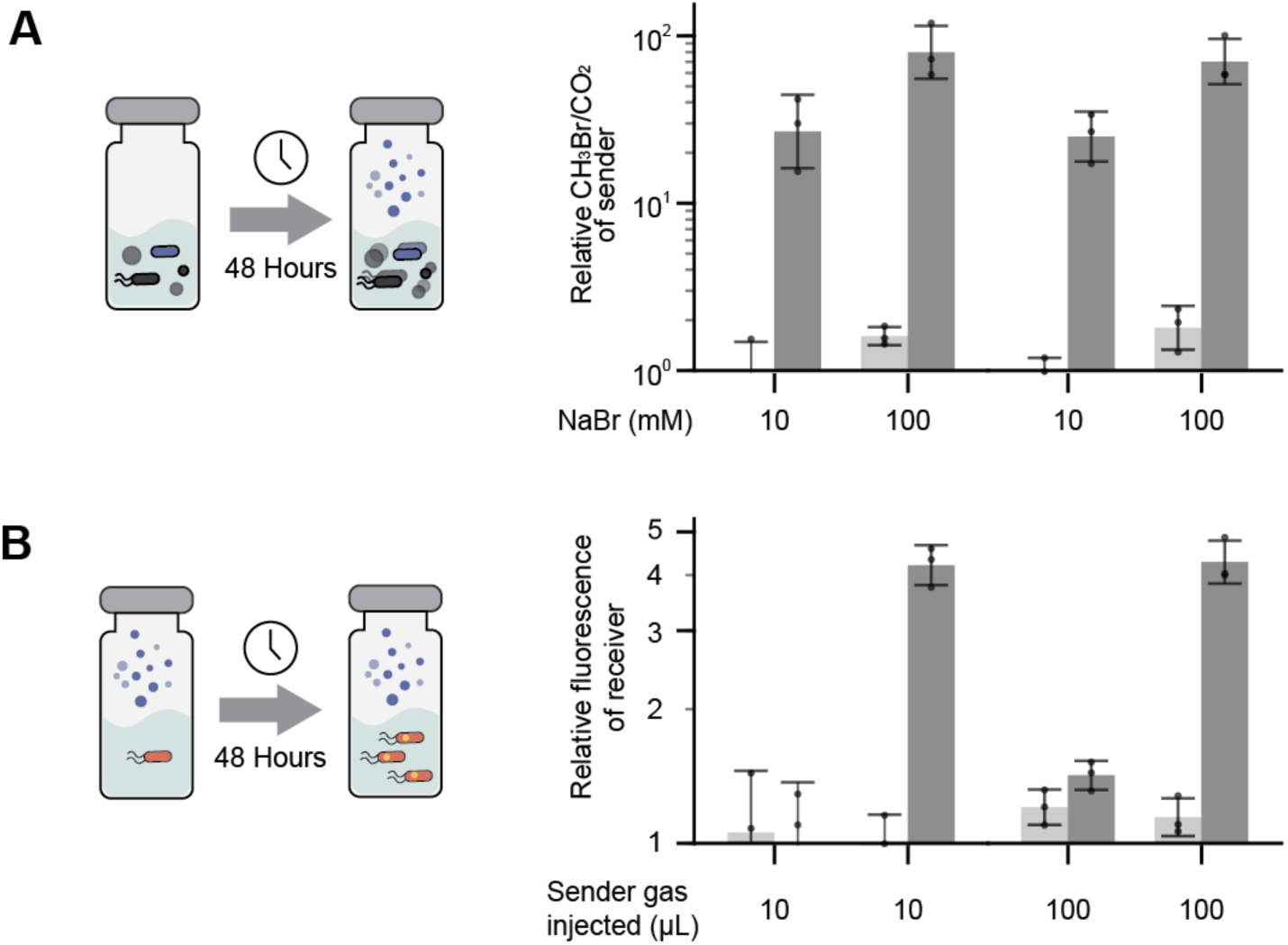
*M. extorquens* can detect CH_3_Br synthesized by a single MSC member. (**A**) An MSC containing *Df, Ea, Rh, St,* and either native *Vb* or *Vb-mht* was incubated for 48 hours in R2A liquid medium supplemented with either 10 or 100 mM NaBr at 30°C. Consortia containing *Vb*-mht produced more CH_3_Br than consortia containing native *Vb* across all conditions tested (*p < 0.05; two-tailed, unpaired, t-test). (**B**) Two different volumes of headspace gas (10 or 100 µL) were transferred from these cultures to *M. extorquens* CM4 cultures programmed to produce YFP upon detecting CH_3_Br as shown in Figure 4a. *M. extorquens* CM4 fluorescence was measured two days after gas transfer. Receiver cells incubated with gas from consortia containing *Vb*-mht grown in 100 mM NaBr presented significantly higher fluorescence than *cells* receiving the same amount of headspace gas from sender consortia containing native *Vb* (*p < 0.01; two-tailed, unpaired, t-test). Bars represent the mean from 3 biological replicates, while error bars represent ±1 standard deviation.

### CH_3_Br cell-cell signaling in soil

The findings that MHT-expressing microbes can generate sufficient headspace CH_3_Br in small GC-MS vials suggested that methyl halide cell-cell signaling could be achieved from a CH_3_Br sender microbe within a bulk soil and CH_3_Br receiver microbe in the headspace of the soil. To test this idea, we designed a simple, easy-to-build soil habitat using commercially available conical tubes (Figure 6). These habitats were designed to allow for facile studies of cell-cell signaling in a closed system containing soil columns up to 10 cm in length where sender cells can be injected at the bottom and receiver cells can be placed in the headspace. To test whether interspecies cell-cell signaling can be achieved in these soil habitats, we performed experiments where sender cells (*E. coli* or *Ec*-mht) in liquid culture were placed at the bottom of the habitat (Figure 6A), and receiver cells (*M. extorquens* CM4) were placed at the top of the habitat. Following a two-day incubation in a closed habitat, the optical density and fluorescence of the receiver cells were measured (Figure S8). These measurements revealed that the whole cell fluorescence was significantly higher in habitats containing *Ec*-mht compared with those containing native *E. coli* (Figure 6B). In addition, when a receiver strain was used that lacked a CH_3_Br-inducible promoter, the fluorescence signal arising from an *Ec*-mht sender was abolished. These results show how a simple soil habitat can be built and used to monitor gas-mediated cell-cell signaling. To investigate whether CH_3_Br-mediated cell-cell signaling could be achieved between a *Vb*-mht (sender) in soil and *M. extorquens* (receiver) in the headspace, we evaluated signaling in a soil habitat. These experiments were performed using soil hydrated to different water holding capacities (6.5, 15, and 33%) using MIDV1 containing 100 mM NaBr and an MSC containing either *Vb*-mht or *Vb*. No antibiotics were included in the soil. After 72 hours, the optical density and fluorescence of the receiver cells were measured (Figure S9). We then compared the relative fluorescence from receiver cells from habitats containing a *Vb*-mht and native *Vb*. Within habitats containing *Vb*-mht, the whole cell fluorescence of the receiver was significantly higher when the soil was hydrated to 15 and 33% water holding capacity (Figure 6C). These results show that a gas-reporting soil microbes can be detected using *M. extorquens* CM4 in the headspace of soil habitats.

**Figure 6.**
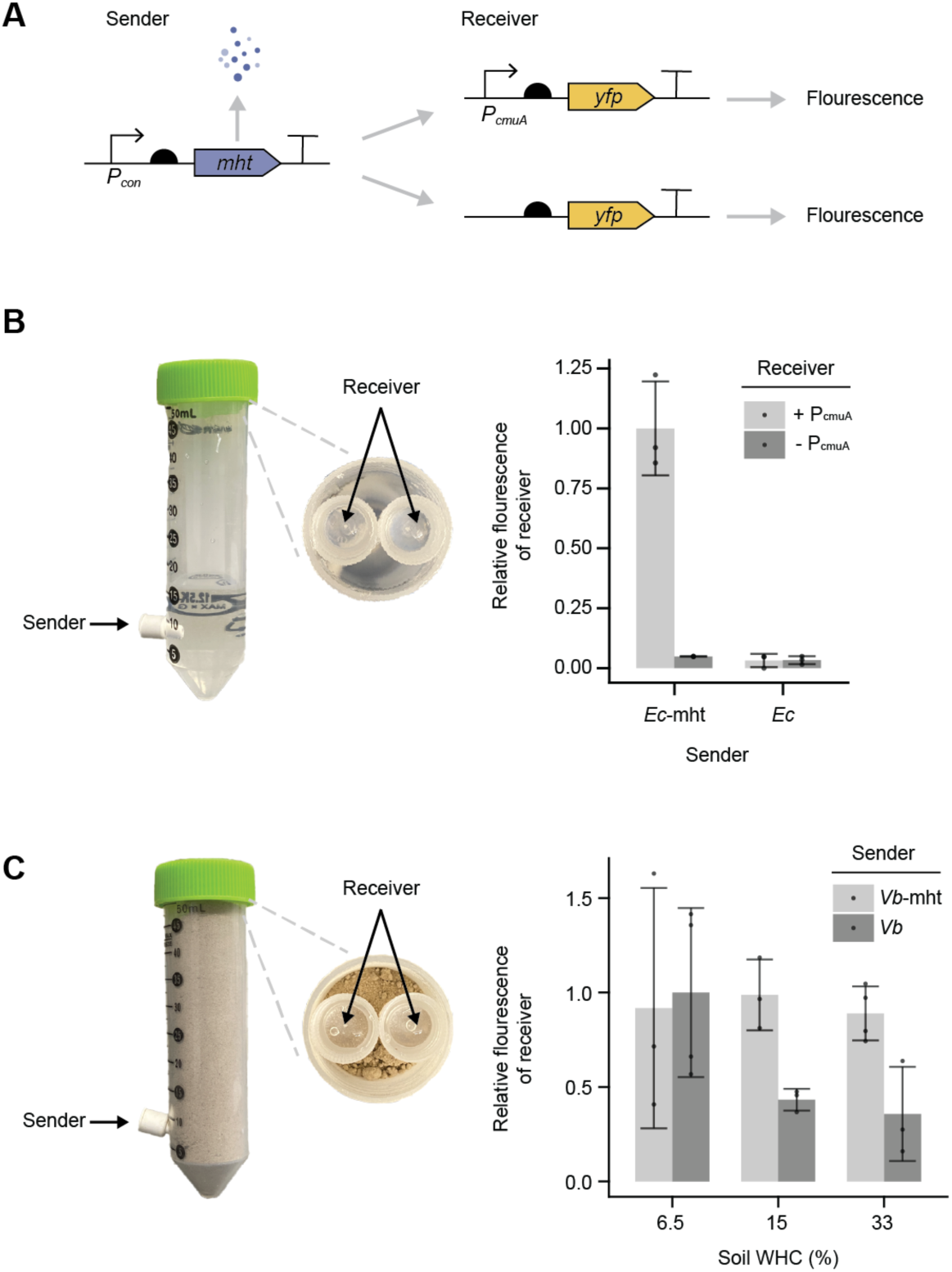
Cell-cell signaling in a synthetic soil habitat. (**A**) Genetic circuits used to program CH_3_Br production in the sender strain and CH_3_Br detection using *M. extorquens* CM4 receiver strain. As a negative control in the receiver, we used a vector that lacked a promoter for the reporter. Commercial conical tubes to study cell-cell signaling using sender cells in (**B**) liquid culture and (**C**) soil. For experiments using liquid medium in the habitat, the relative fluorescence of receiver cells in the headspace of habitats was monitored following incubation of *Ec-mht* or native *E. coli* MG1655 in the base of the habitats. The receiver strain presented a higher signal than the negative strain when the sender expressed MHT (*p<0.05; two-tailed, unpaired, t-test). For the soil experiments, the relative fluorescence of receiver cells in the headspace of habitats was measured following incubation in habitats filled to capacity with soil and hydrated to different water holding capacities (WHC) using medium containing *Vb* or *Vb-mht*. The receiver strain had a higher YFP signal with the *Vb*-mht as a sender compared to native *Vb* in soil hydrated to 15% and 33% WHC (*p<0.05; two-tailed, unpaired, t-test). All incubations were performed for 72 hours in closed soil habitats. Each bar represents the mean of 3 biological replicates and error bars represent ±1 standard deviation.

### Implications

Our studies show for the first time that gas reporters are compatible with monitoring growth and conditional gene expression in a MSC made up of interacting microbes that can be stored, revived, and shared.^11^ Our analysis of a putative hyperosmotic responsive promoter shows how MSC regulatory elements can be characterized *in vitro* by varying the concentrations of osmolytes, which alters osmotic pressure, and *in situ* where the matrix properties (*e.g.*, soil particle size and hydration) also contribute to the water potential due to the capillary and adsorptive forces between the liquid and the solid phase of soils.^40^ Osmotic responsive promoters are typically characterized in liquid cultures,^41^ and it remains unclear how osmolytes and soil work together to regulate the activities of soil microbe promoters through changes in the total soil water potential, which is regulated by the sum of osmotic matric potential in addition to gravitational and pressure potential.^40^ Since wet-dry cycles affect both osmotic and matric potentials in soil, a better understanding of such dynamic regulation will be important for understanding bursts of respiration and nitrification that occur when dry soils are rewetted, such as the Birch effect.^28, 42^ In the future, gas reporters can be combined with -omics measurements to better understand the effects of different combinations of osmotic and matric potentials on gene expression *in situ*.

The finding that methyl halide cell-cell signaling can be achieved between different soil microbes in a cm-scale soil habitat extends the volatile signals that can be used to program information flow in a community beyond acetaldehyde and hydrogen sulfide.^43, 44^ In future studies, it will be interesting to explore whether such volatile cell-cell signaling can be used to create synthetic networks of information flow that report on the bioavailability of organic pollutants in soil and sediments,^45^ the state of bioremediation of organic and inorganic pollutants,^46, 47^ and soil health.^48^ To extend these studies, it will be important to tune this signaling so that it functions in other habitats developed for bulk soil studies, such as EcoFABs and rhizotrons.^49, 50^ To be useful for studies *in situ*, it will be important to explore how this type of cell-cell signaling can be used to overcome challenges at the soil-air interface where volatile chemical signals are massively diluted by the atmosphere, thereby limiting detection using analytical methods.^18^ This signaling approach could be built upon to create a volatile-to-nonvolatile signal converter at the soil-air interface to extend the lifetime of any sensed information generated by an indicator gas reporter.

## Methods

### Materials

Enzymes for molecular biology (Phanta Max Super-Fidelity DNA polymerase, T4 Ligase, BsaI, and DpnI) were from Vazyme Biotech and New England Biolabs. Primers for cloning were from Integrated DNA Technologies. DNA amplicons were purified using Zymoclean Gel DNA Recovery Kit (Zymo Research), and plasmids were purified from cells using the QIAprep Spin Miniprep Kit (Qiagen). Sodium bromide was from MilliPoreSigma, methyl bromide standards were from Restek, and all other growth media components and chemicals were from MilliPoreSigma, Fisher, Restek, Research Products International, and VWR. Gas tight 2 mL glass vials (AR0-37L0-13) and verex seals (AR0-5760-13) used to crimp vials were from Phenomenex.

### Strains

All molecular biology was performed using either *Escherichia coli* DH10B or NEB Turbo, while indicator gas studies were performed using *E. coli* MG1655 or *E. coli*-mht, a strain that constitutively expresses *B. maritima* MHT from the chromosome.^18^ The five MSC members were from a natural soil consortium sampled from Warden silt loam in Prosser, WA.^7, 11^ The conjugation of plasmids into *V. beijingensis* was achieved using *E. coli* MFD*pir*.^51^ *M extorquens* CM4 was used as a receiver strain for synthetic cell-cell signaling, since it contains a promoter activated by methyl halides.^22^ To establish the antibiotic sensitivity of each MSC member (Table S1), strains were streaked on R2A-agar containing a range of *Vb* carbenicillin, chloramphenicol, kanamycin, spectinomycin, streptomycin, and tetracycline concentrations. Plates were grown at 19°C until colonies have a diameter of at least 0.5 mm were observed on plates lacking antibiotics, which ranged from 2 to 5 days. At this time point, the lowest antibiotic concentration that suppressed growth compared to the control plate was established.

### Plasmids

Table S2 lists all plasmids used. Methyl halide biosensing was achieved using a *M. extorquens* CM4 transformed with a plasmid (pME8266) that uses the *P_cmuA_* promoter to regulate YFP expression.^22^ A promoterless version of this vector (pLM-sYFP2) was used as a control to evaluate *M. extorquens* CM4 autofluorescence. A broad host range plasmid for constitutive gas production (pPK114) was created that uses the cumate-inducible promoter (*P_cym_*) to control the transcription of MHT;^52^ it contains a pBBR1 origin and kanR. Translation initiation was achieved using the ribosome binding sites BBa_B0034 and BBa_B0030, respectively. The plasmid for conditionally expressing MHT using the CDS.21 promoter (pJWK020) was created by replacing the promoter in pPK114 with 150 bp of DNA that is upstream of *Vb* CDS.21;^27^ this was achieved using Golden Gate Assembly.^53^ This sequence was from the Genome Taxanomy Database. All vectors were sequence verified.

### Conjugation

To perform conjugation, *E. coli* MFD*pir* was first grown in LB medium supplemented with 0.3 mM diaminopimelic acid (DAP) and kanamycin (50 µg/mL) overnight. In parallel, the receiver strains were grown at 30°C in R2A medium to an OD of at least 0.5. Cells from these cultures were washed with sterile phosphate buffered saline (PBS), mixed at 1:1 ratio based on their OD (100 µL), placed on top of LB-agar medium containing 0.3 mM DAP, and incubated at 30°C. After 48 hours, the cell mixture was resuspended in PBS, washed for 3 times to remove DAP, and spread on LB-agar medium containing kanamycin (50 µg/mL) but lacking DAP. After incubating at 30°C for 48 hours, cells were were streaked out onto a new LB-agar plate containing kanamycin to obtain single colonies of transconjugants. *Vb* were chosen based on their yellow morphology and the gain of antibiotic resistance.

### MSC gas production

To assess gas production from individual MSC members, cells derived from single colonies were grown in R2A liquid medium at 30°C, while shaking at 250 rpm. After 36 to 48 hours, cultures were diluted to an OD_600_ of 0.05 in fresh R2A medium (1 mL) containing 10 mM or 100 mM NaBr within glass vials (2 mL), which were immediately crimp sealed. After incubation for 48 hours at 30°C while shaking at 250 rpm, the optical density and headspace gas were measured.

### Headspace gas analysis

A gas chromatograph (GC) mass spectrometer (MS) consisting of an Agilent 8890 GC and a 5977B MS was used for headspace gas analysis. An Agilent 7693A autosampler fitted with a 100 µL gastight syringe was used to inject headspace gas (50 µL) into a DB-VRX capillary column (20 m, 0.18 mm ID, 1 µm film; Agilent). Experiments were performed using a 55:1 split ratio. The oven temperature was initially held at 45°C for 84 seconds before ramping to 60°C at 0.6°C/seconds and then holding at 60°C for 9 seconds. MS analysis was performed using selected ion monitoring mode for CH_3_Br (m/z = 93.9 and 95.9) and CO_2_ (m/z = 44 and 45). Peak area was quantified using the Agilent MassHunter Workstation Quantitative Analysis software. To construct a CH_3_Br standard curve, liquid CH_3_Br standards (Restek, 30253) were serially diluted into M63 medium (1 mL) in gas tight vials (2 mL). After allowing equilibration for 4 hours, the CH_3_Br signals in the headspace of these samples were quantified using GC-MS. A standard curve was generated (Figure S10), which was used to convert GC-MS peak areas into absolute concentrations of CH_3_Br in the liquid medium.

### Growth medium

Overnight cultures of cells grown in R2A (*Vb*) and LB (*E. coli*) were washed by centrifuging at 3000 g (5 min) and resuspending in either R2A, M63, or MIDV1, a diluted minimal medium.^25^ In each condition, cells were diluted to an OD of 0.05. Cells were then grown at 30°C while shaking at 250 rpm for 48 hours prior to headspace gas analysis using GC-MS. R2A medium (per Liter) contained 0.5 g yeast extract, 0.5 g proteose peptone No. 3, 0.5 g casamino acids, 0.5 g glucose, 0.5 g soluble starch, 0.3 g sodium pyruvate, 0.3 g K_2_HPO_4_, and 0.05 g MgSO_4_ᐧ7H_2_O.^54^ M63 medium (per Liter) was made by mixing 50 mL of 20% (w/v) glucose, 1 mL of 1M MgSO_4_, 100 µL of 0.1% (w/v) thiamine hydrochloride, 5 mL of 10% (w/v) casamino acids, and 200 mL of M63 salt stock. M63 salt contained 75 mM ammonium sulfate, 0.5 M potassium phosphate monobasic, and 10 µM ferrous sulfate. This salt stock was adjusted to pH 7 and autoclaved prior to use.^19^ MIDV1 medium (per liter) was made by mixing 50 mL of 20% (w/v) glucose, 1 mL of 1M MgSO_4_, 100 µL of 0.1% (w/v) thiamine hydrochloride, 10 mL of 10% (w/v) casamino acids, and 12.5 mL of M63 salt stock.^25^ NaBr was added at the indicated levels.

### Soil vial incubations

All soil studies were performed using a previously characterized Texas Alfisol,^20^ a Oxyaquic Glossudalf. After sampling, the soil was immediately dried by baking at 60°C and then sieved (2 mm) prior to storage at room temperature in sealed containers. To prepare *E. coli* and *V. beijingensis* for soil measurements, they were grown to stationary phase in LB (37°C) and R2A (30°C), respectively, while shaking at 250 rpm. In cases where cells were transformed with vectors, the growth medium contained kanamycin (50 μg/mL). Cells were then washed 3 times and resuspended in MIDV1 medium (1 mL) containing 100 mM NaBr. After diluting cultures to an OD_600_ of 0.05 using MIDV1, this solution was used to hydrate unautoclaved soil (1 gram) to 32% or 64% of the field capacity using GC-MS vials (2 mL). After crimping, vials were incubated statically at 30°C for 48 hours prior to headspace gas analysis.

### Osmotic stress measurements

To characterize the *Vb* CDS.21 promoter (Pvb0021), individual *Vb* colonies transformed with either pJWK020 or empty vector (pPK048) were grown overnight in LB medium containing kanamycin (10 µg/mL). Stationary phase cultures were pelleted and washed three times using MIDV1. Cells were then diluted to an OD_600_ of 0.05 in MIDV1 medium (1 mL) supplemented with 20 mM NaBr varying sucrose concentrations (0, 100, 200, 300, 400, 500, and 600 mM) within glass vials (2 mL). These sucrose concentrations in MIDV1 yielded solutions with osmotic potentials of -319, -567, -815, -1063, -1311, -1559, and -1807 kPa. Vials were immediately crimped, and cells were then grown at 30°C while shaking at 250 rpm. After 48 hours, headspace gas was measured, and the optical density was measured. Experiments in soil used a similar protocol, with the exception that washed cells were added to 1g of an artificial soil (Q3a) that mimics the texture of clay.^25^ With experiments performed in artificial soils, incubations were static. For soils, the potential reported represents the sum of the osmotic and matric potential; the latter was previously measured for this soil.^25^

### CH_3_Br biosensor calibration

To generate cultures that produce CH_3_Br, *E. coli*-mht were grown in LB (5 mL) for 16 hours at 37°C while shaking at 250 rpm, cells were pelleted, pellets were resuspended in M63 containing 100 mM NaBr,^20^ and the washed cells were diluted to a final OD_600_ of 0.05 (1 mL) in 2 mL vials. Vials were crimped and incubated at 37°C for 24 hours while shaking at 250 rpm. The headspace gas was measured using a GC-MS, and a gastight syringe (Hamilton, 1700 series) was used to transfer different volumes (1, 2, 5, 10, 50 and 100 µL) of headspace gas to a second crimped vial which contained *M. extorquens* CM4 (1 mL). To generate CH_3_Br biosensor cultures, *M. extorquens* CM4 transformed with pME8266 were grown on LB-agar containing kanamycin (50 µg/mL) to obtain single colonies, which were used to inoculate nsLB medium (5 mL) containing 50 µg/mL kanamycin, a growth medium that is identical to LB except it omits sodium chloride. After growing cells to mid log phase at 30°C while shaking at 250 rpm in 14 mL Falcon tubes (Fisher Scientific, 352059), cells were diluted to an OD_600_ = 0.1 (1 mL) in 2 mL vials and capped. The headspace gas from donor cultures that had already been grown for 24 hours were immediately injected into vials containing *M. extorquens* cultures. Following gas injection, these latter vials were incubated at 30°C for 48 hours while shaking at 250 rpm. Fluorescence and absorbance (OD600) were then analyzed.

Immediately prior to gas transfer, the amount of headspace gas in the *E. coli* cultures was analyzed to estimate the amount injected. The CH_3_Br counts from GC-MS were converted to molar concentrations using the CH_3_Br standard curve. To calculate the CH_3_Br concentration in receiver vials, we first calculated the total amount of CH_3_Br present in the sender vials by converting the concentrations obtained from the x-axis of the standard curve (Figure S10) to absolute amounts by multiplying the concentration with the volume of aqueous solution (in dm^3^). The total amount of CH_3_Br present in each vial can be expressed as [CH_3_Br]_total_ = C_a_/1000 + C_g_/1000, where *C_a_* and *C_g_* are the concentrations of CH_3_Br in the aqueous phase and the gas-phase of the vials, respectively. Based on Henry’s law, *H^CC^* = C_a_/C_g_ and *H^CC^ = H^CP^* x *RT,* where *H^CP^* is the Henry’s solubility constant. The *H^CP^*value for CH_3_Br has been previously reported.^55^ Hence we calculate *H^CC^* given the *H^CP^* values reported (1.7×10^−3^ mol/m^3^ Pa). Using this value of *H^CP^*, one obtains a *H^CC^*value of 4.280. We then used the *H^CC^* value to obtain C_a_ = 4.28·C_g_. To obtain the absolute amount of CH3Br in the head space of sender vials, we calculated the product of *C_g_* and the total volume of headspace gas (0.001 dm^3^). This value was used to calculate the amount of CH_3_Br transferred to receiver vials. The amount of CH_3_Br transferred to the receiver vial was then used to calculate C_a_ and C_g_, which allowed us to calculate the CH_3_Br concentration encountered by the *M. extorquens* receiver cells growing in solution.

### MSC gas-transfer experiments

Soils containing an MSC used a mixture of each community member, each at OD600 = 0.05. To generate the MSC, single colonies of each strain were grown in R2A medium (5 mL) for 40 hours (*St* and *Vb*) or 64 hours (*Df, Ea, and Rh*) at 30°C while shaking at 250 rpm. Cultures containing *Vb* transformed with vectors additionally contained kanamycin (50 µg/mL). Cells were washed in R2A medium containing 10 or 100 mM NaBr and resuspended to an OD600 = 0.05 in fresh medium. To generate the MSC, cultures from each microbe (200 µL) were mixed together in a 2 mL vial, such that they contained a native MSC (*Df, Ea, Rh, St, and Vb) or a MSC where the Vb* is transformed with the vector that constitutively produces MHT. Prior to crimping vials, a fraction of each MSC sample was frozen to enable analysis of microbial diversity before incubation. The remaining samples were crimped and incubated at 30°C for 48 hours while shaking at 250 rpm. The headspace gas from these culture mixtures were analyzed GC-MS, and a gastight syringe was then used to transfer defined volumes (10 and 100 μL) to the headspace of crimped vials containing *M. extorquens* at an OD_600_ = 0.1 (1 mL). *M. extorquens* CM4 signals were measured after a 48 hour incubation.

### Fluorescence analysis

To quantify YFP in *M. extorquens* CM4 following sensor incubations, cultures (200 µL) were transferred to 96-well flat bottom black microplates (Corning, 3603). A Tecan Spark or a Tecan M1000 plate reader was then used to measure absorbance at 600 nm and whole cell fluorescence (λ_ex_ = 485 nm; λ_em_ = 516 nm, bandwidth = 5 nm). Measurements were also performed using growth medium, and these values were subtracted from whole cell data prior to calculating the fluorescence to absorbance ratios. The ratiometric signals were then normalized against the maximum value to obtain relative fluorescence values.

### Conical habitats assembly

To construct conical habitats,50 mL conical tubes (VWR, 89039-656) were modified by drilling a hole (3/16 inch diameter) at ∼5 mm above the bottom of the tube. To make air tight, a rubber septum was inserted with an 8 mm outer diameter (MilliporeSigma, Z553913). To sterilize habitats, they were soaked in 70% ethanol for 2 hours and left to air-dry in a biosafety cabinet. To perform signaling studies in soil within these conical habitats, they were filled to the 50 mL line with the Texas Alfisol using a sterile spatula and cryotube lids were placed on the soil surface; cryotube lids (ThermoFisher Scientific, 5000-0020) were glued inside the tubes exactly 10 mm from the upper rim using Clear Gorilla® Glue (Target, 77657594) for liquid cultures studies. Habitats were then capped, and the soil was hydrated to the indicated water holding capacity (WHC) by injecting cells resuspended in MIDV1 into the rubber septum using an 18 gauge syringe needle (Thomas Scientific, 305196). In the case of *E. coli*, which were used for initial liquid culture experiments, 6.4 ✕ 10^7^ CFUs of *Ec*-mht or native E. coli in M63 (16 mL) were injected into the habitats using an 18-gauge syringe needle (Thomas Scientific, 305196). In the case of *Vb*, the volume injected varied, but the total number of cells was held constant at 5 x10^8^ CFUs. After adding cells to the bottom of the habitat, the conical tubes were uncapped, and receiver cells were added to the cryotube lids (500 μL) at an OD600 of 0.1 in nsLB containing kanamycin (50 µg/mL). Habitats were capped, sealed with parafilm, and incubated at room temperature (20°C) for 72 hours.

### Statistical analysis

A minimum of three biological replicates were performed in each experiment. *p-values* determined by performing independent two-tailed t-tests using GraphPad Prism (version 9.3.0). Statistically significant difference was defined by the t-tests returning a *p*-value of less than 0.05.

## Abbreviations

(Df): Dyadobacter fermentans
(Ea): Ensifer adhaerens
(Ec): Escherichia coli
(GC-MS): Gas chromatography mass spectrometry
(MHT): Methyl Halide Transferase
(MSC): Model Soil Consortium
(OD): Optical Density
(Rh): Rhodococcus sp003130705
(St): Streptomyces sp001905665
(Vb): Variovorax beijingensis
(YFP): Yellow Fluorescent Protein
(WHC): Water Holding Capacity

## Author Contributions

Contributions are noted in alphabetical order.

Conceptualization: *CAM*, *JJS, and KSH*

Methodology: *CAM, JJS, JK, LCL, and KSH*

Investigation: *JK and LCL*

Visualization: *JJS, JK, and LCL*

Project administration: *CAM*, *JJS, and KSH*

Supervision: *CAM*, *KSH*, and *JJS*

Writing – original draft: *JJS, JK, and LCL*

Writing – review & editing: *CAM, JJS, JK, LCL, KSH, and RM*

## ACKNOWLEDGEMENTS

This research was supported by Defense Advanced Research Projects Agency contract HR0011-19-2-0019 and a subcontract from the US Department of Energy, Office of Science, through the Genomic Science Program, Office of Biological and Environmental Research, under FWP 78814 at PNNL (to CAM and JJS). PNNL is a multi-program national laboratory operated by Battelle for the DOE under Contract DE-AC05-76RLO 1830.

## REFERENCES

(1) Keesstra, S., Geissen, V., Mosse, K., Piiranen, S., Scudiero, E., Leistra, M., and Van Schaik, L. (2012) Soil as a filter for groundwater quality. Current Opinion in Environmental Sustainability 4, 507–516.

(2) Voigt, C. A. (2020) Synthetic biology 2020-2030: six commercially-available products that are changing our world. Nat Commun 11, 6379.

(3) Jansson, J. K., and Hofmockel, K. S. (2020) Soil microbiomes and climate change. Nat Rev Microbiol 18, 35–46.

(4) Bardgett, R. D., and van der Putten, W. H. (2014) Belowground biodiversity and ecosystem functioning. Nature 515, 505–511.

(5) Wagg, C., Bender, S. F., Widmer, F., and van der Heijden, M. G. A. (2014) Soil biodiversity and soil community composition determine ecosystem multifunctionality. Proc Natl Acad Sci U S A 111, 5266–5270.

(6) Hultman, J., Waldrop, M. P., Mackelprang, R., David, M. M., McFarland, J., Blazewicz, S. J., Harden, J., Turetsky, M. R., McGuire, A. D., Shah, M. B., VerBerkmoes, N. C., Lee, L. H., Mavrommatis, K., and Jansson, J. K. (2015) Multi-omics of permafrost, active layer and thermokarst bog soil microbiomes. Nature 521, 208–212.

(7) McClure, R., Farris, Y., Danczak, R., Nelson, W., Song, H.-S., Kessell, A., Lee, J.-Y., Couvillion, S., Henry, C., Jansson, J. K., and Hofmockel, K. S. (2022) Interaction Networks Are Driven by Community-Responsive Phenotypes in a Chitin-Degrading Consortium of Soil Microbes. mSystems 7, e0037222.

(8) Burmeister, A., and Grünberger, A. (2020) Microfluidic cultivation and analysis tools for interaction studies of microbial co-cultures. Curr Opin Biotechnol 62, 106–115.

(9) Prosser, J. I. (2015) Dispersing misconceptions and identifying opportunities for the use of “omics” in soil microbial ecology. Nat Rev Microbiol 13, 439–446.

(10) Jansson, J. K., and Hofmockel, K. S. (2018) The soil microbiome-from metagenomics to metaphenomics. Curr Opin Microbiol 43, 162–168.

(11) McClure, R., Naylor, D., Farris, Y., Davison, M., Fansler, S. J., Hofmockel, K. S., and Jansson, J. K. (2020) Development and Analysis of a Stable, Reduced Complexity Model Soil Microbiome. Front Microbiol 11, 1987.

(12) Geddes, B. A., Paramasivan, P., Joffrin, A., Thompson, A. L., Christensen, K., Jorrin, B., Brett, P., Conway, S. J., Oldroyd, G. E. D., and Poole, P. S. (2019) Engineering transkingdom signalling in plants to control gene expression in rhizosphere bacteria. Nat Commun 10, 3430.

(13) Ryu, M.-H., Zhang, J., Toth, T., Khokhani, D., Geddes, B. A., Mus, F., Garcia-Costas, A., Peters, J. W., Poole, P. S., Ané, J.-M., and Voigt, C. A. (2020) Control of nitrogen fixation in bacteria that associate with cereals. Nat Microbiol 5, 314– 330.

(14) Chen, S., Chang, C., Deng, Y., An, S., Dong, Y. H., Zhou, J., Hu, M., Zhong, G., and Zhang, L.-H. (2014) Fenpropathrin biodegradation pathway in Bacillus sp. DG-02 and its potential for bioremediation of pyrethroid-contaminated soils. J Agric Food Chem 62, 2147–2157.

(15) Russo, F., Ceci, A., Pinzari, F., Siciliano, A., Guida, M., Malusà, E., Tartanus, M., Miszczak, A., Maggi, O., and Persiani, A. M. (2019) Bioremediation of Dichlorodiphenyltrichloroethane (DDT)-Contaminated Agricultural Soils: Potential of Two Autochthonous Saprotrophic Fungal Strains. Appl Environ Microbiol 85, e01720–19.

(16) Jiménez-Díaz, V., Pedroza-Rodríguez, A. M., Ramos-Monroy, O., and Castillo- Carvajal, L. C. (2022) Synthetic Biology: A New Era in Hydrocarbon Bioremediation. Processes 10, 712.

(17) Kolosova, E. M., Sutormin, O. S., Shpedt, A. A., Stepanova, L. V., and Kratasyuk, V. A. (2022) Bioluminescent-Inhibition-Based Biosensor for Full-Profile Soil Contamination Assessment. Biosensors (Basel) 12, 353.

(18) Cheng, H.-Y., Masiello, C. A., Bennett, G. N., and Silberg, J. J. (2016) Volatile Gas Production by Methyl Halide Transferase: An In Situ Reporter Of Microbial Gene Expression In Soil. Environ Sci Technol 50, 8750–8759.

(19) Cheng, H.-Y., Masiello, C. A., Del Valle, I., Gao, X., Bennett, G. N., and Silberg, J. J. (2018) Ratiometric Gas Reporting: A Nondisruptive Approach To Monitor Gene Expression in Soils. ACS Synth Biol 7, 903–911.

(20) Fulk, E. M., Gao, X., Lu, L. C., Redeker, K. R., Masiello, C. A., and Silberg, J. J. (2022) Nondestructive Chemical Sensing within Bulk Soil Using 1000 Biosensors Per Gram of Matrix. ACS Synth Biol 11, 2372–2383.

(21) Del Valle, I., Webster, T. M., Cheng, H.-Y., Thies, J. E., Kessler, A., Miller, M. K., Ball, Z. T., MacKenzie, K. R., Masiello, C. A., Silberg, J. J., and Lehmann, J. (2020) Soil organic matter attenuates the efficacy of flavonoid-based plant-microbe communication. Sci Adv 6, eaax8254.

(22) Farhan Ul Haque, M., Nadalig, T., Bringel, F., Schaller, H., and Vuilleumier, S. (2013) Fluorescence-based bacterial bioreporter for specific detection of methyl halide emissions in the environment. Appl Environ Microbiol 79, 6561–6567.

(23) Redeker, K. R., and Cicerone, R. J. (2004) Environmental controls over methyl halide emissions from rice paddies: ENVIRONMENTAL CONTROLS OVER MeX FLUXES FROM RICE. Global Biogeochem. Cycles 18, n/a-n/a.

(24) Wuosmaa, A. M., and Hager, L. P. (1990) Methyl chloride transferase: a carbocation route for biosynthesis of halometabolites. Science 249, 160–162.

(25) Del Valle, I., Gao, X., Ghezzehei, T. A., Silberg, J. J., and Masiello, C. A. (2022) Artificial Soils Reveal Individual Factor Controls on Microbial Processes. mSystems 7, e0030122.

(26) Satola, B., Wübbeler, J. H., and Steinbüchel, A. (2013) Metabolic characteristics of the species Variovorax paradoxus. Appl Microbiol Biotechnol 97, 541–560.

(27) Gao, J.-L., Sun, Y.-C., Xue, J., Sun, P., Yan, H., Khan, M. S., Wang, L.-W., Zhang, X., and Sun, J.-G. (2020) Variovorax beijingensis sp. nov., a novel plant-associated bacterial species with plant growth-promoting potential isolated from different geographic regions of Beijing, China. Syst Appl Microbiol 43, 126135.

(28) Schimel, J. P. (2018) Life in Dry Soils: Effects of Drought on Soil Microbial Communities and Processes. Annu. Rev. Ecol. Evol. Syst. 49, 409–432.

(29) Yim, H. H., and Villarejo, M. (1992) osmY, a new hyperosmotically inducible gene, encodes a periplasmic protein in Escherichia coli. J Bacteriol 174, 3637–3644.

(30) Hoffmann, T., Wensing, A., Brosius, M., Steil, L., Völker, U., and Bremer, E. (2013) Osmotic control of opuA expression in Bacillus subtilis and its modulation in response to intracellular glycine betaine and proline pools. J Bacteriol 195, 510– 522.

(31) Kakumanu, M. L., and Williams, M. A. (2014) Osmolyte dynamics and microbial communities vary in response to osmotic more than matric water deficit gradients in two soils. Soil Biology and Biochemistry 79, 14–24.

(32) Chowdhury, N., Marschner, P., and Burns, R. G. (2011) Soil microbial activity and community composition: Impact of changes in matric and osmotic potential. Soil Biology and Biochemistry 43, 1229–1236.

(33) Del Valle, I., Fulk, E. M., Kalvapalle, P., Silberg, J. J., Masiello, C. A., and Stadler, L. B. (2020) Translating New Synthetic Biology Advances for Biosensing Into the Earth and Environmental Sciences. Front Microbiol 11, 618373.

(34) Patel, K. F., Fansler, S. J., Campbell, T. P., Bond-Lamberty, B., Smith, A. P., RoyChowdhury, T., McCue, L. A., Varga, T., and Bailey, V. L. (2021) Soil texture and environmental conditions influence the biogeochemical responses of soils to drought and flooding. Commun Earth Environ 2, 127.

(35) Bardgett, R. D., and Caruso, T. (2020) Soil microbial community responses to climate extremes: resistance, resilience and transitions to alternative states. Phil. Trans. R. Soc. B 375, 20190112.

(36) Garg, A., Bordoloi, S., Ganesan, S. P., Sekharan, S., and Sahoo, L. (2020) A relook into plant wilting: observational evidence based on unsaturated soil–plant-photosynthesis interaction. Sci Rep 10, 22064.

(37) Papendick, R. I., and Campbell, G. S. (2015) Theory and Measurement of Water Potential, in SSSA Special Publications (Parr, J. F., Gardner, W. R., and Elliott, L. F., Eds.), pp 1–22. Soil Science Society of America, Madison, WI, USA.

(38) Studer, A., McAnulla, C., Büchele, R., Leisinger, T., and Vuilleumier, S. (2002) Chloromethane-induced genes define a third C1 utilization pathway in Methylobacterium chloromethanicum CM4. J Bacteriol 184, 3476–3484.

(39) Chaignaud, P., Maucourt, B., Weiman, M., Alberti, A., Kolb, S., Cruveiller, S., Vuilleumier, S., and Bringel, F. (2017) Genomic and Transcriptomic Analysis of Growth-Supporting Dehalogenation of Chlorinated Methanes in Methylobacterium. Front Microbiol 8, 1600.

(40) Wraith, J., and Or, D. (2001) Soil Water Content and Water Potential Relationships, in Soil Physics Companion (Warrick, A., Ed.), pp 49–84. CRC Press.

(41) Bianchi, A. A., and Baneyx, F. (1999) Hyperosmotic shock induces the sigma32 and sigmaE stress regulons of Escherichia coli. Mol Microbiol 34, 1029–1038.

(42) Birch, H. F. (1958) The effect of soil drying on humus decomposition and nitrogen availability. Plant Soil 10, 9–31.

(43) Weber, W., Daoud-El Baba, M., and Fussenegger, M. (2007) Synthetic ecosystems based on airborne inter- and intrakingdom communication. Proc. Natl. Acad. Sci. U.S.A. 104, 10435–10440.

(44) Liu, H., Fan, K., Li, H., Wang, Q., Yang, Y., Li, K., Xia, Y., and Xun, L. (2019) Synthetic Gene Circuits Enable *Escherichia coli* To Use Endogenous H _2_ S as a Signaling Molecule for Quorum Sensing. ACS Synth. Biol. 8, 2113–2120.

(45) Zhang, X., Li, B., Schillereff, D. N., Chiverrell, R. C., Tefsen, B., and Wells, M. (2022) Whole-cell biosensors for determination of bioavailable pollutants in soils and sediments: Theory and practice. Sci Total Environ 811, 152178.

(46) Eze, M. O., Thiel, V., Hose, G. C., George, S. C., and Daniel, R. (2022) Bacteria-plant interactions synergistically enhance biodegradation of diesel fuel hydrocarbons. Commun Earth Environ 3, 192.

(47) Tay, P. K. R., Nguyen, P. Q., and Joshi, N. S. (2017) A Synthetic Circuit for Mercury Bioremediation Using Self-Assembling Functional Amyloids. ACS Synth Biol 6, 1841–1850.

(48) Lehmann, J., Bossio, D. A., Kögel-Knabner, I., and Rillig, M. C. (2020) The concept and future prospects of soil health. Nat Rev Earth Environ 1, 544–553.

(49) Sasse, J., Kant, J., Cole, B. J., Klein, A. P., Arsova, B., Schlaepfer, P., Gao, J., Lewald, K., Zhalnina, K., Kosina, S., Bowen, B. P., Treen, D., Vogel, J., Visel, A., Watt, M., Dangl, J. L., and Northen, T. R. (2019) Multilab EcoFAB study shows highly reproducible physiology and depletion of soil metabolites by a model grass. New Phytol 222, 1149–1160.

(50) Huck, M. G., and Taylor, H. M. (1982) The Rhizotron as a Tool for Root Research, in Advances in Agronomy, pp 1–35. Elsevier.

(51) Ferrières, L., Hémery, G., Nham, T., Guérout, A.-M., Mazel, D., Beloin, C., and Ghigo, J.-M. (2010) Silent mischief: bacteriophage Mu insertions contaminate products of Escherichia coli random mutagenesis performed using suicidal transposon delivery plasmids mobilized by broad-host-range RP4 conjugative machinery. J Bacteriol 192, 6418–6427.

(52) Eaton, R. W. (1997) p-Cymene catabolic pathway in Pseudomonas putida F1: cloning and characterization of DNA encoding conversion of p-cymene to p-cumate. J Bacteriol 179, 3171–3180.

(53) Engler, C., Kandzia, R., and Marillonnet, S. (2008) A one pot, one step, precision cloning method with high throughput capability. PLoS One 3, e3647.

(54) Reasoner, D. J., and Geldreich, E. E. (1985) A new medium for the enumeration and subculture of bacteria from potable water. Appl Environ Microbiol 49, 1–7.

(55) Sander, R. (2015) Compilation of Henry’s law constants (version 4.0) for water as solvent. Atmos. Chem. Phys. 15, 4399–4981.

