## Supporting Information for "Using methyl bromide for interspecies cell-cell signaling and as a reporter in a model soil consortium"

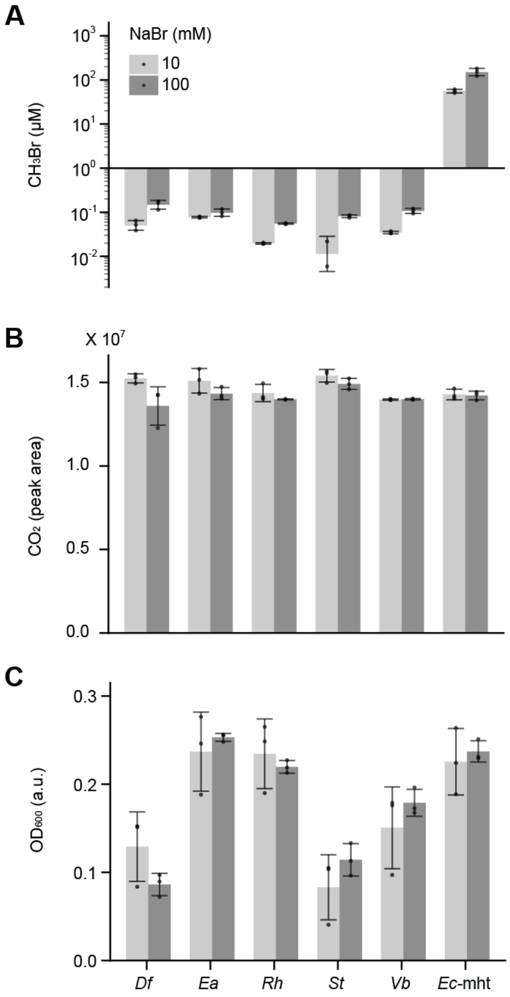


**Figure S1. Indicator gas, carbon dioxide, and optical density for individual MSC members.** Individual MSC members were grown in R2A medium (1 mL) supplemented with 10 mM (light gray) or 100 mM NaBr (dark gray) while shaking at 250 rpm in sealed vials. Experiments were also performed using an *E. coli* strain (*Ec*-mht) that constitutively expresses MHT from the chromosome. After incubating 48 hours, GC-MS was used tro measure (**A**) $CH_3Br$ and (**B**) carbon dioxide in the headspace. (**C**) Following the headspace measurement, optical density was measured. Error bars represent ±1 standard deviation from three biological replicates.

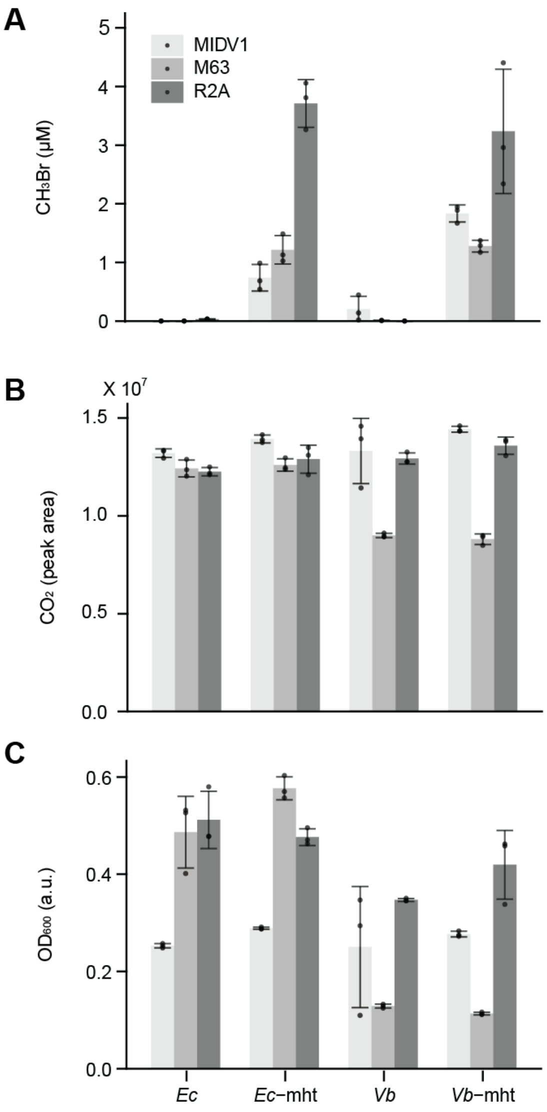


**Figure S2. Effect of liquid medium on indicator gas production, respiration, and growth. (A)** Methyl bromide and **(B)** carbon dioxide produced by *E. coli* (*Ec*) and *V. beijingensis* (*Vb*) grown in different liquid medium (R2A, M63, and MIDV1) supplemented with 100 mM NaBr. Gas production was evaluated from cells alone and cells transformed with a vector that constitutively express *B. maritima* MHT (*Ec*-mht and *Vb*-mht). (**C**) The optical density of cultures at the end of each incubation. Error bars represent ±1 standard deviation from three biological replicates.

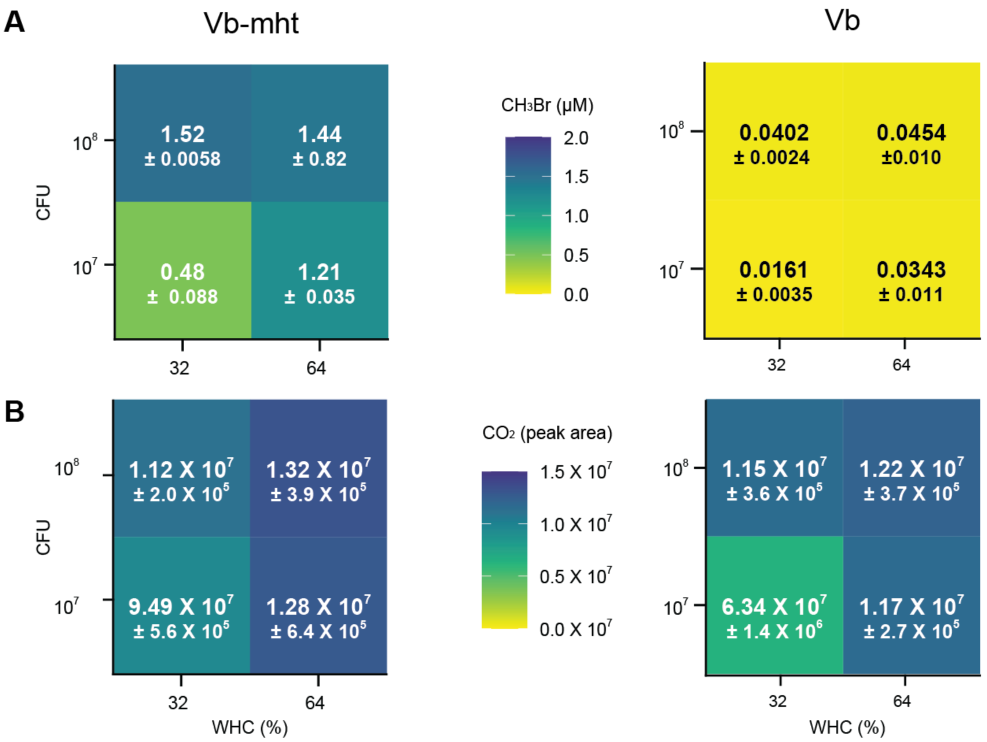


**Figure S3. $CH_3Br$ and $CO_2$ in the headspace of soils containing *V. beijingensis*.** (A) The $CH_3Br$ signal in the headspace of Texas Alfisol soil inoculated with *V. beijingensis* containing (*left*) and lacking (*right*) a vector that constitutively expresses an MHT (pPK114). (**B**) The $CO_2$ signal in the headspace of soils inoculated with *V. beijingensis* containing (*left*) and lacking (*right*) a vector that constitutively expresses an MHT. Values shown represent the mean from three biological replicates ±1 standard deviation.

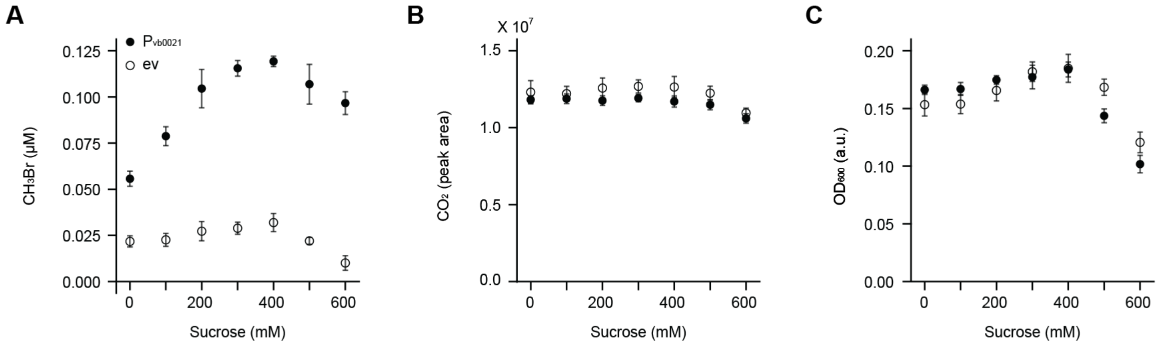


**Figure S4. Effect of water potential on a *V. beijingensis* promoter monitored using gas reporting.** *V. beijingensis* cells transformed with either a putative osmolarity biosensing plasmid (pJWK020) or a empty vector (pPK048) were incubated in 1 mL MIDV1 (20 mM NaBr) supplemented with varying concentrations of sucrose at 30 °C, while shaking at 250 rpm, for 48 hours. (**A**) $CH_3Br$ and (B) carbon dioxide were measured using GC-MS. (**C**) Following the gas measurements, optical density ($OD_{600}$) was analyzed. Each point plotted represents the arithmetic mean of 5 biological replicates and error bars represent ±1 standard deviation.

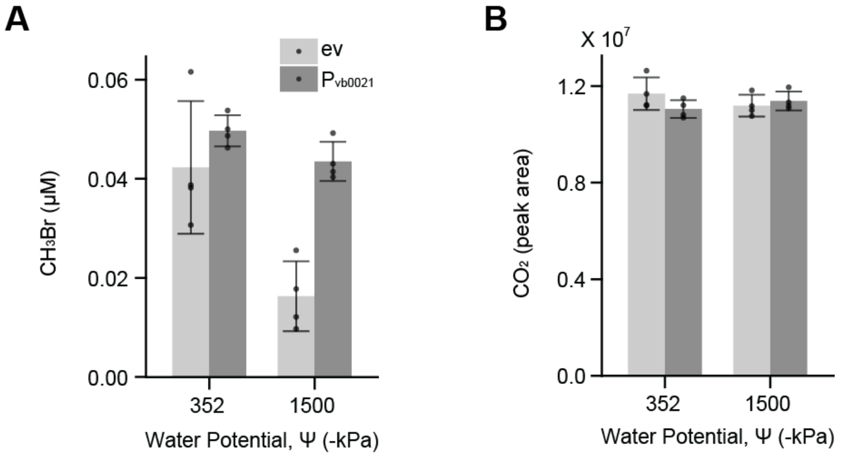


**Figure S5. Respiration and gas production in soil.** *V. beijingensis* cells transformed with a plasmid (pJWK020) that uses an uncharacterized promoter, Pvb0021 to regulate MHT expression was incubated in a clay-mimic artificial soil hydrated to 100% field capacity with MIDV1 minimal medium (20 mM NaBr) and incubated at 30 °C for 48 hours. **(A)** $CH_3Br$ and **(B)** respiration (were measured using GC-MS. *Vb* transformed with pJWK020 produces similar amount of indicator gas as *Vb* transformed with empty vector (ev) at -352 kPa (**p > 0.01, two-tailed, ujpaired t-test). *Vb* transformed with pJWK020 produces significantly higher indicator gas than *Vb* transformed with empty vector (ev) when sucrose is added to increase the pressure to -1500 kPa (p > 0.05, two-tailed, unpaired t-test). Each bar plotted represents the arithmetic mean of 4 biological replicates and error bars represent ±1 standard deviation.

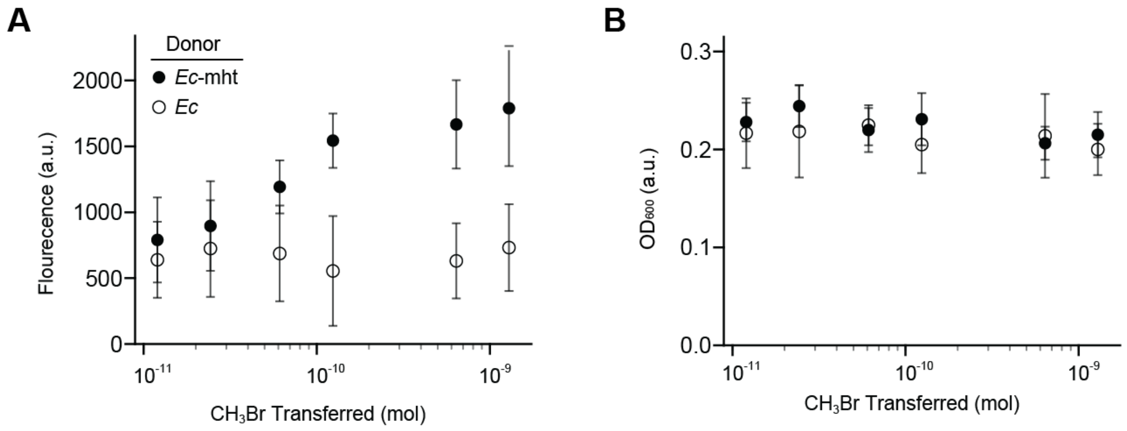


**Figure S6. Fluorescent protein signal from *M. extorquens* receivers following headspace gas injections. (A)** YFP fluorescence and **(B)** optical density of *M. extorquens* CM4 following injection of gas from closed cultures containing different *E. coli*. Two different volumes (10, 100 μL) of headspace gas from a sealed vial containing *E. coli* expressing an MHT were injected into vials containing a *M. extorquens* CM4 that contains the YFP gene under control of the $P_{cmuA}$ promoter (closed circle). As a negative control, native *E. coli* was used as a sender (open circle). Error bars represent ±1 standard deviation from seven biological replicates.

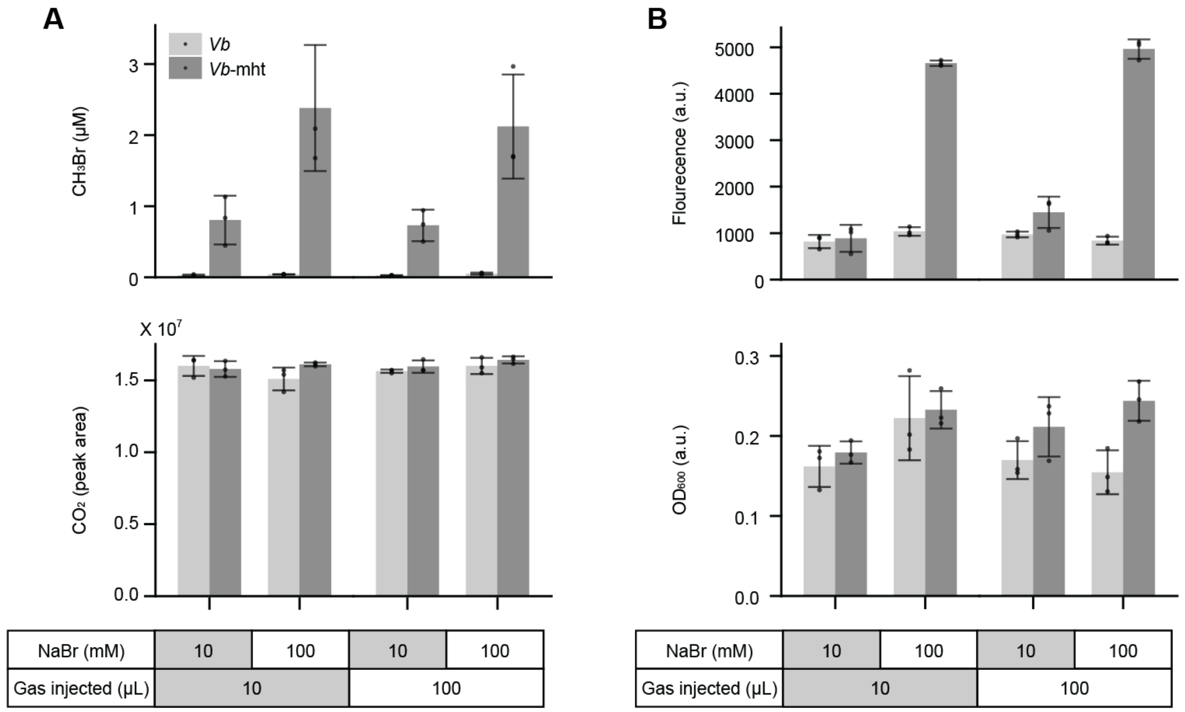


**Figure S7. Sender cell gas production and receiver cell fluorescence following headspace gas transfer. (A)** An MSC containing either native *Vb* (light gray) or *Vb* transformed with a vector that expresses MHT (dark gray), pPK114, was grown in R2A liquid medium supplemented with either 10 or 100 mM NaBr for 48 hours at 30°C. $CH_3Br$ (top) and $CO_2$) (bottom) were measured using GC-MS. A consortium containing *Vb*-mht produced more $CH_3Br$ than consortium containing native *Vb* across all conditions tested (*$p < 0.05$; two-tailed, unpaired, t-test). **(B)** Different volumes (10, 100 µL) of headspace gas from a sealed vial containing MSC with either native *Vb* or *Vb*-mht were injected into vials containing a *M. extorquens* CM4 that contains the YFP gene under control of the $P_{cmuA}$ promoter. Whole cell fluorescence (top) and optical density (bottom) were measured after 48 hours of incubation at 30 °C. Bars represent the mean from 3 biological replicates, while error bars represent ±1 standard deviation.

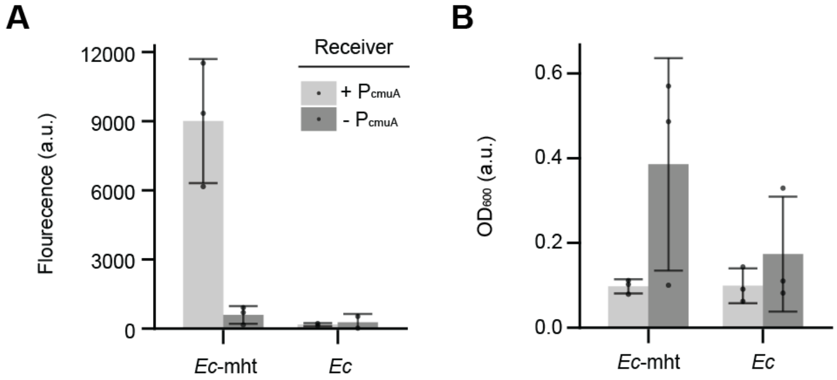


**Figure S8. Cell-cell signaling in a conical habitat containing liquid medium. (A)** Fluorescence and **(B)** optical density of the *M. extorquens* CM4 receiver strain that expresses YFP under control of the $P_{cmuA}$ promoter following incubation in a synthetic conical habitat (50 mL) with *E. coli* that constitutively expresses MHT (*Ec*-mht) or lacks an MHT (*Ec*). As a negative control, *M. extorquens* CM4 was used with a vector that lacks $P_{cmuA}$ promoter. Bars represent the mean from 3 biological replicates, while error bars represent ±1 standard deviation.

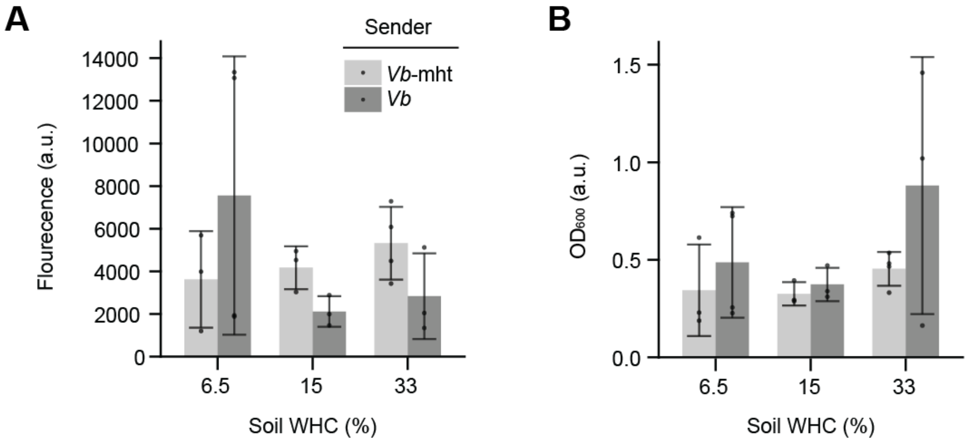


**Figure S9. Cell-cell signaling in a conical habitat containing soil. (A)** Fluorescence and **(B)** optical density of the *M. extorquens* CM4 receiver strain that expresses YFP under control of the $P_{cmuA}$ promoter followingincubation in a synthetic conical habitat with soil hydrated with medium containing a *Vb* strain constitutively express MHT (*Vb*-mht) or native *Vb*. Incubations were performed for 48 hours at room temperature. The conical habitat was filled to capacity with soil and hydrated to different WHCs (6.5, 15, and 33%) with liquid medium containing *Vb.* The receiver cells were incubated on top of the soil. Bars represent the mean from 4 biological replicates, while error bars represent ±1 standard deviation.

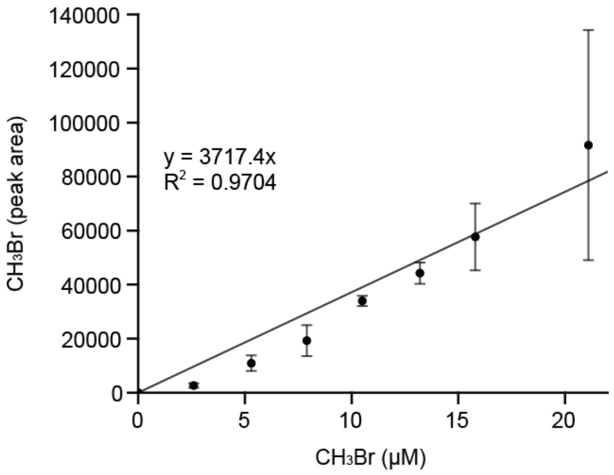


**Figure S10. GC-MS standard curve for methyl bromide.** Different amounts of $CH_3Br$ were added to sealed vials containing M63 medium (1 mL), vials were incubated for 4 hours while shaking at 250 rpm to allow for gas partitioning, and GC-MS was used to measure the headspace gas. Error bars represent ±1 standard deviation from 3 biological replicates. Data was best fit by a linear model (y = 3717.4x, $R^2$ = 0.9704).

**Table S1. Antibiotic susceptibility of different MSC members.** The first column provides the names of each antibiotic and the concentration (µg/mL). Cells were restreaked from the R2A plate lacking antibiotic to an R2A plate containing antibiotic and a second plate lacking antibiotic. Following incubation for two to five days, colonies were observed on all plates lacking antibiotics. If the colonies on plates containing antibiotic were not visible when colonies became visible on the plate lacking antibiotics, it was marked as susceptible.

| | ***St*** | ***Df*** | ***Ea*** | ***Vb*** | ***Rh*** |
|---|---|---|---|---|---|
| TET 2.5 | susceptible | susceptible | susceptible | | susceptible |
| TET 5 | susceptible | susceptible | susceptible | susceptible | susceptible |
| TET 10 | susceptible | susceptible | susceptible | susceptible | susceptible |
| TET 20 | susceptible | susceptible | susceptible | susceptible | susceptible |
| KAN 5 | susceptible | | | susceptible | |
| KAN 10 | susceptible | | | susceptible | |
| KAN 20 | susceptible | | | susceptible | susceptible |
| KAN 50 | susceptible | | | susceptible | susceptible |
| CHL 5 | | | | | susceptible |
| CHL 10 | | | | | susceptible |
| CHL 20 | susceptible | | | | susceptible |
| CHL 50 | susceptible | | | | susceptible |
| CARB 10 | | | | | |
| CARB 20 | | | | | |
| CARB 50 | | | | | |
| CARB 100 | | | | | |
| SPEC 10 | | | | | |
| SPEC 20 | | | | | |
| SPEC 50 | susceptible | | susceptible | | |
| SPEC 100 | susceptible | | susceptible | susceptible | |
| STREP 10 | susceptible | | | | susceptible |
| STREP 20 | susceptible | | | | susceptible |
| STREP 50 | susceptible | | | susceptible | susceptible |
| STREP 100 | susceptible | | | susceptible | susceptible |

**Table S2. Plasmids used in this study.**

| Plasmid | Description | Promoter | Origin of replication | Antibiotic Resistance Marker |
|---|---|---|---|---|
| pJWK020 | hyperosmotic response regulated MHT expression | *Pvb.cds.21* | pBBR1 | Kan |
| pLM-sYFP2 | promoterless YFP | *n/a* | colE1 | Kan |
| pME8266 | Methyl halide-regulated YFP expression | $P_{cmuA}$ | colE1 | Kan |
| pPK114 | Constitutive MHT expression | $P_{cym}$ | pBBR1 | Kan |
| pPK048 | Constitutive RFP expression, negative control for $CH_3X$ producing plasmid | $P_{cym}$ | pBBR1 | Kan |
